# High Sox2 expression predicts taste lineage competency of lingual progenitors in vitro

**DOI:** 10.1101/2022.06.30.498302

**Authors:** Lauren A. Shechtman, Jennifer K. Scott, Eric D. Larson, Trevor J. Isner, Bryan J. Johnson, Dany Gaillard, Peter J. Dempsey, Linda A. Barlow

## Abstract

Taste buds on the tongue are collections of taste receptor cells (TRCs) that detect sweet, sour, salty, umami and bitter stimuli. Like non-taste lingual epithelium, TRCs are renewed from basal keratinocytes, many of which express the transcription factor SOX2. Genetic lineage tracing has shown SOX2+ lingual progenitors give rise to both taste and non-taste lingual epithelium in the posterior circumvallate taste papilla (CVP) of mice. However, SOX2 is variably expressed among CVP cells suggesting that their progenitor potential may vary. Using transcriptome analysis and organoid technology, we show highly expressing SOX2+ cells are taste-competent progenitors that give rise to organoids comprising both TRCs and lingual epithelium, while organoids derived from low-expressing SOX2+ progenitors are composed entirely of non-taste cells. Hedgehog and WNT/ß-catenin are required for taste homeostasis in adult mice, but only WNT/ß-catenin promotes TRC differentiation *in vitro* and does so only in organoids derived from higher SOX2+ taste lineage-competent progenitors.

## Introduction

Gustation is mediated by taste buds in specialized structures on the tongue: in rodents, each fungiform papilla (FFP) on the anterior tongue houses a single bud, while posterior foliate and circumvallate papillae (CVP), house hundreds of taste buds. Each taste bud contains ∼60 taste receptor cells (TRCs) categorized as: type I glial-like support cells, type II cells that detect sweet, bitter, or umami stimuli, and type III cells that respond to sour and some salty stimuli (Roper and Chaudhari, 2017). TRCs and surrounding non-taste epithelium are continuously replenished by basal progenitor cells; these give rise directly to non-taste lingual epithelium, and to taste-fated daughter cells that enter buds, transiently express Sonic Hedgehog (SHH) and differentiate into each of the TRC types (Finger and Barlow, 2021). To date, progenitors expressing LGR5 (leucine-rich repeat-containing G-protein coupled receptor 5) (Takeda et al., 2013; Yee et al., 2013), GLI1 (Liu et al., 2013), cytokeratin (KRT) 14 (Okubo et al., 2009), KRT5 (Gaillard et al., 2015), and SOX2 (SRY-related HMG box family)(Ohmoto et al., 2017) are known to give rise to TRCs and non-taste epithelium in mice, but their potentially distinct roles in taste epithelial renewal are unexplored. In murine CVP, LGR5+ progenitors also function as taste stem cells *in vitro*, as isolated LGR5+ cells generate lingual organoids housing cycling progenitors, non-taste epithelium and TRCs (Ren et al., 2014). However, the potential of SOX2+ progenitors to generate TRC-replete organoids *in vitro* has not been explored.

SOX2 is a key regulator of homeostasis in many adult epithelia (Arnold et al., 2011; Novak et al., 2020), whose function often depends on expression level (Hagey and Muhr, 2014; Que et al., 2009; Sarkar and Hochedlinger, 2013). SOX2 is expressed by most basal keratinocytes within taste epithelium (Castillo-Azofeifa et al., 2018; Suzuki, 2008), and long-term lineage tracing of SOX2+ cells labels taste and non-taste epithelia in mice (Ohmoto *et al*., 2017). However, lingual SOX2 expression is highly variable with highest expression in a subset of TRCs and in progenitors immediately adjacent to buds. These findings suggest high SOX2-expressing progenitors in lingual epithelium replenish taste buds, whereas low SOX2-expressing basal cells may not. Here, we use organoid technology to test if SOX2 expression level predicts the ability of isolated progenitors to give rise to TRCs *in vitro* and show only higher expressing SOX2+ progenitors are competent to produce TRC-replete organoids.

## Results

### LGR5 and SOX2 expression partially overlap in mouse CVP epithelium

We first assessed SOX2 immunofluorescence (IF) in *Lgr5*^*EGFP*^ mice (Barker et al., 2007). As reported (Ohmoto *et al*., 2017; Suzuki, 2008), SOX2-IF is strong in some TRCs and basal cells adjacent to buds but is low in cells in the deep CVP trench (**Fig 1A-A’’**). As previously reported (Yee *et al*., 2013), LGR5-GFP is bright in cells in the deep CVP, and less robust along the trench walls (**Fig 1A-A”**). *Sox2* mRNA is highly expressed by cells in and around taste buds, but low in the deep trench, consistent with SOX2-IF. However, *Lgr5* expression levels are similar throughout the CVP epithelium, in contrast to pattern of LGR5-driven GFP which is strongest deep in the trench (Compare **Fig 1A** to **1B**). This discrepancy may be due to perdurance of GFP (Arnone et al., 2004), a phenomenon likely enhanced in the LGR5+ slower cycling cells of the deep trench (Yee *et al*., 2013). Nonetheless, CVP basal keratinocytes co-express SOX2 and LGR5, albeit with differing relative intensities.

**Figure 1.**
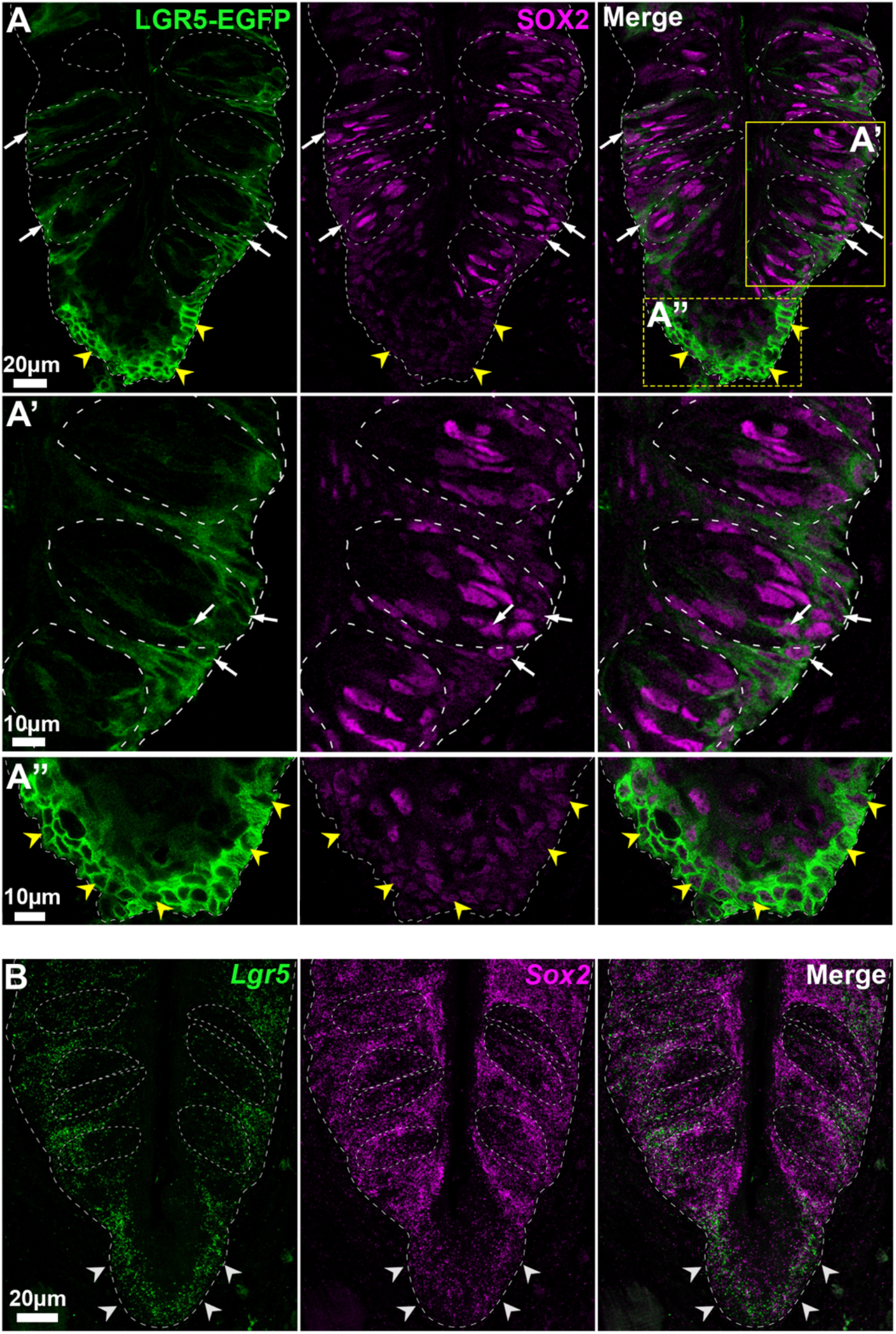
LGR5 and SOX2 expression overlap in mouse CVP. **A-A”** SOX2-IF (magenta) is robust in and around taste buds (dashed circles), dimmer in LGR5-GFP+ cells (green) in trench wall epithelium (white arrows), and low in deep CVP epithelium where LGR5-GFP is highest (yellow arrowheads). **B** *Lgr5* mRNA (green) is comparably expressed by cells at the basement membrane (dashed line) deep in the CVP (white arrowheads) and associated with taste buds (dashed circles), while *Sox2* mRNA expression (magenta) is high in and around buds but low in the deep CVP epithelium. Images are optical sections in **A-A”** and maximum projections of confocal z-stacks in **B**.

### SOX2+ progenitors have limited potential to generate TRC-replete organoids

SOX2+ and LGR5+ cells generate TRCs and non-taste epithelium *in vivo*, but only LGR5+ cells have been shown to produce TRC-replete organoids (see (Barlow, 2021). Thus, we employed *Sox2*^*GFP*^ mice where GFP reliably reports SOX2 expression in taste epithelium (Castillo-Azofeifa *et al*., 2018; Okubo et al., 2006) to determine if isolated SOX2+ progenitors generate TRC-containing organoids. SOX2-GFP+ and LGR5-GFP+ cells were isolated and cultured as described (Shechtman et al., 2021) (**Fig 2A**). Quantitative PCR (qPCR) revealed that mRNA for markers of type I (*Entpd2, Kcnj1*)(Bartel et al., 2006; Dvoryanchikov et al., 2009), type II (*Gnat3, Pou2f3*) (Boughter et al., 1997; Matsumoto et al., 2011) and type III (*Car4, Pkd2l1*) (Kataoka et al., 2008; Wilson et al., 2017) TRCs was expressed by SOX2-derived organoids (SOX2 organoids) but at significantly lower levels than LGR5-derived organoids (LGR5 organoids) (**Fig 2B**). Via IF, we found most LGR5 organoids contained type I (NTPDase2+), type II (GUSTDUCIN+), and type III (CAR4+) TRCs as expected, while most SOX2 organoids lacked TRCs (**Fig 2C**). Further, LGR5 organoids had significantly more TRCs of each type (>10 taste cells/organoid), whereas the few SOX2 organoids with TRCs had fewer cells per organoid (**Fig 2D**). Sparse NTPDase2+ and GUSTDUCIN+ cells in SOX2 organoids appeared poorly differentiated, lacking the typical fusiform morphology of type I and II cells in LGR5 organoids (**Fig 2C**). Nonetheless, a small number of SOX2 organoids were TRC-replete, suggesting a subset of SOX2+ progenitors are taste competent.

**Figure 2.**
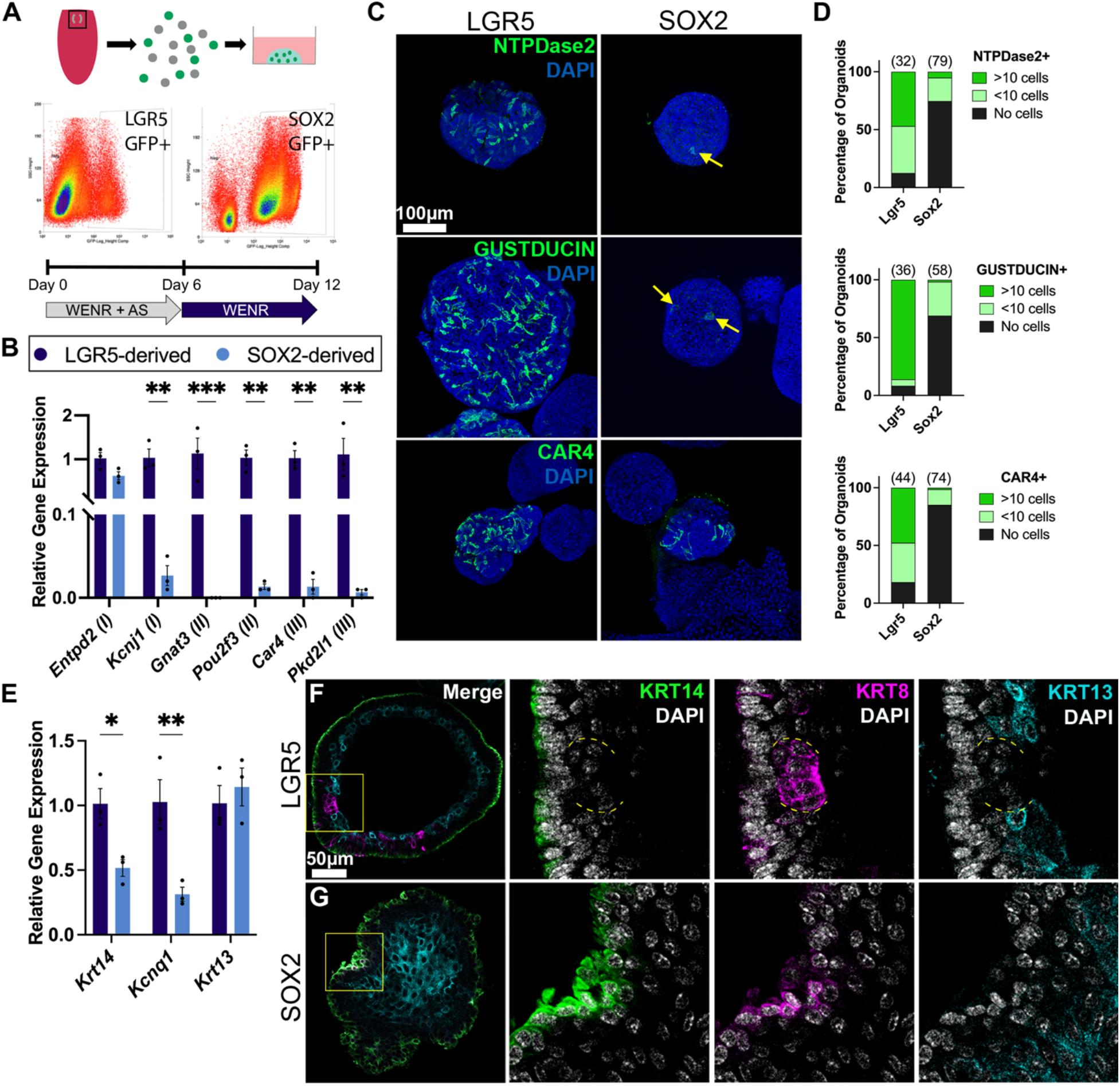
SOX2+ progenitors have limited potential to generate TRC-replete organoids. **A** Procedure to generate organoids from CVP progenitors from *Lgr5*^*EGFP*^ or *Sox2*^*EGFP*^ mice. **B** Type I (*Entpd2, Kcnj1*), II (*Gnat3, Pou2f3*), and III (*Car4, Pkd2l1*) TRC marker expression is significantly lower in SOX2 vs LGR5 organoids. **C, D** Most LGR5 organoids contain type I (NTPDase2+ green), II (GUSTDUCIN+ green) and III (CAR4 green) TRCs, but most SOX2 organoids do not. TRCs in SOX2 organoids lack conventional taste cell morphology (yellow arrows). Images are maximum projections of confocal z-stacks of whole organoids. In **D**, (n) = organoid number analyzed per condition. **E** LGR5 organoids have higher *Kcnq1* (TRCs) and *Krt14 (*progenitors) but similar *Krt13* (non-taste epithelium) compared to SOX2 organoids. **F** In LGR5 organoids, KRT14+ cells (green) are basal/external and KRT8+ TRCs (magenta) (yellow dash outline) are internal and surrounded by KRT13+ non-taste cells (cyan). **G** SOX2 organoids contain mostly KRT13+ cells (cyan); KRT8 (magenta) and KRT14 (green) are often co-expressed. Images are optical sections of immunostained organoids. DAPI nuclear counterstain blue in **C**, white in **F, G**. Scale bar in **F** also for **G**. For **B** and **E** mean ± SEM, n = 3 biological replicates, 2-way ANOVA, * p<0.05, ** p<0.01, *** p<0.001.

As SOX2+ progenitors generate few organoids with TRCs, we posited the remainder comprised primarily non-taste epithelium. Consistent with this, *Kcnq1* (general TRC marker) (Wang et al., 2009) and *Krt14* (cycling progenitors) were significantly lower in SOX2 vs LGR5 organoids, while *Krt13* (non-taste marker) was comparable between organoid types (**Fig 2E**). In LGR5 organoids, KRT14+ progenitors made up the organoid exterior, while differentiated TRCs (KRT8+) and non-taste cells (KRT13+) were situated internally, reflecting the basal/external - apical/internal organization of lingual organoids (**Fig 2F**) (Ren *et al*., 2014). KRT14+ cells were also external in SOX2 organoids, but internally these organoids comprised mainly KRT13+ non-taste cells (**Fig 2G**). Additionally, KRT8 and KRT14 were frequently co-expressed by cells at the periphery of SOX2 organoids (**Fig 2G**). In adult CVP as progenitors produce new TRCs, KRT14 is downregulated and KRT8 upregulated, so that immature cells within taste buds are transiently KRT8+ and KRT14+ (Asano-Miyoshi et al., 2008). In SOX2 organoids, these KRT8+/14+ cells may therefore be immature TRCs (**Fig 2G**). In sum, while some SOX2 organoids are taste competent, most are composed of non-taste epithelium with little TRC differentiation.

### Progenitors with differential SOX2 expression are transcriptionally distinct

Because TRC production is limited in SOX2 organoids, and higher SOX2 expressing cells are associated with taste buds *in vivo*, we hypothesized that higher SOX2+ populations would have gene profiles consistent with taste lineage production. To explore this, SOX2^High^, SOX2^HiMed^, SOX2^MedLow^, and SOX2^Low^ cells were collected via FACS from *Sox2*^*GFP*^ CVP epithelium and processed for bulk RNA sequencing and analysis. Relative expression of *Sox2* and *Gfp* confirmed the accuracy of sorted SOX2-GFP+ brightness bins (**Fig 3A, File S1**). Gene expression profiles across biological replicates within brightness bins were highly consistent; transcriptomes of SOX2^HiMed^ and SOX2^MedLow^ cells were similar, and SOX2^High^ and SOX2^Low^ cells most distinct (**Fig 3B**).

**Figure 3.**
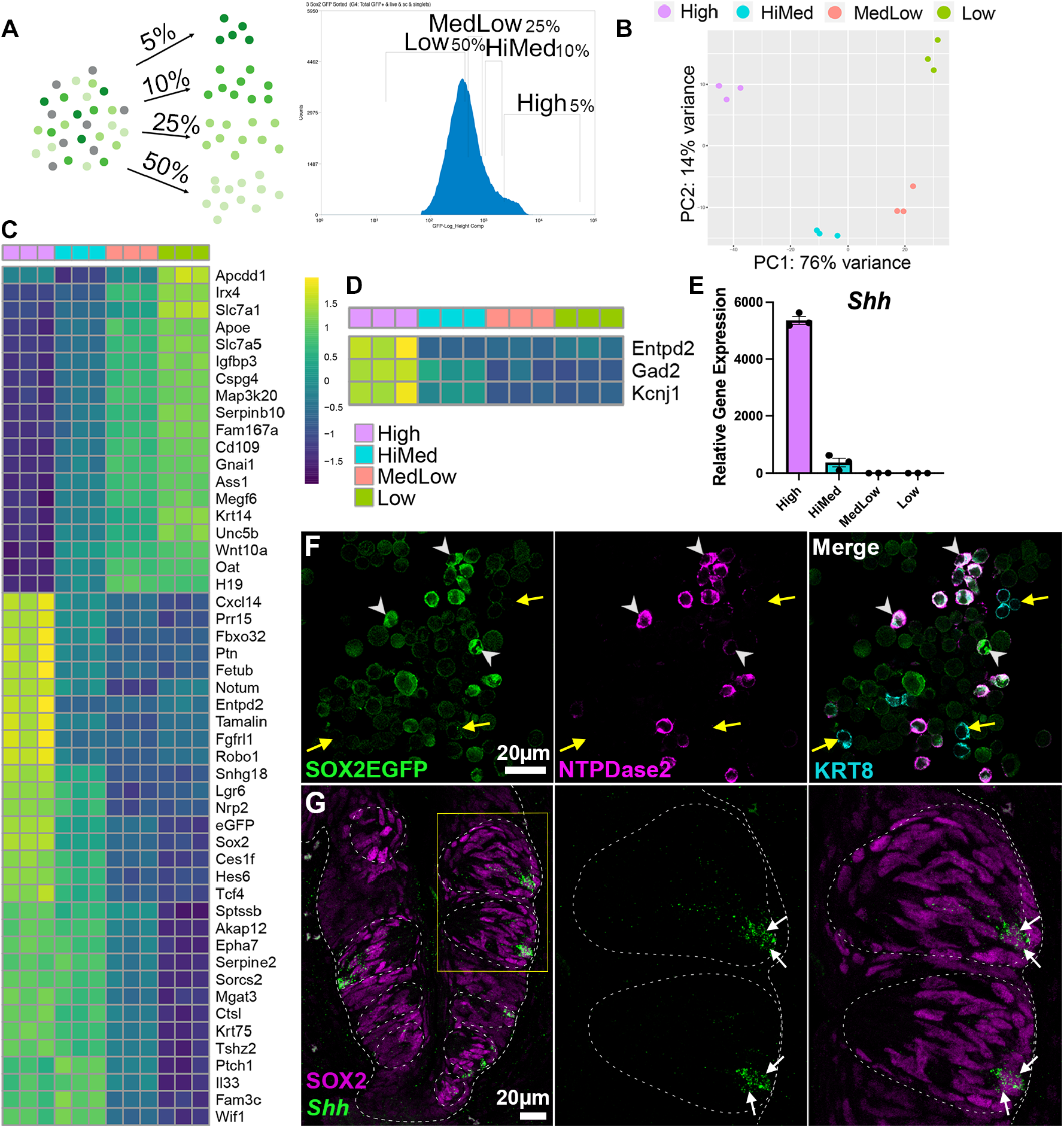
Cells with differential SOX2 expression are transcriptionally distinct. **A** SOX2-GFP+ progenitors were separated via FACS into 4 brightness bins. **B** Biological replicates of each SOX2-GFP population cluster appropriately via principal component analysis. **C** Top 50 genes (by FDR-adjusted p value) differentially expressed across the 4 populations identified by likelihood ratio test. **D, E** Type I TRC markers and *Shh* are enriched in SOX2^High^ cells. **F** In a dispersed cell preparation of CVP epithelium from *Sox2*^EGFP^ mice, KRT8+ (cyan) taste cells that express high SOX2-GFP (green) are NTPDase2+ (magenta, white arrowheads), while KRT8+ taste cells lacking SOX2-GFP are NTPDase2^neg^ (yellow arrows). Image is an optical section. **G** *Shh* (green) is expressed in SOX2-IF cells (magenta, white arrows) in CVP taste buds (dashed circles). Image is a confocal z-stack projection.

In addition to the top 50 differentially expressed genes (FDR adjusted p < 0.05) (**Fig 3C**), type I markers were enriched in SOX2^High^ cells (**Fig 3D**). Published reports suggest SOX2 is expressed by type I cells in CVP taste buds (Suzuki, 2008; Takeda *et al*., 2013); however, identifying individual type I TRCs in tissue sections is problematic (see (Miura et al., 2014). NTPDase2-IF of dispersed CVP cells from *Sox2*^*GFP*^ mice revealed many SOX2-GFP^Bright^ cells were NTPDase2/KRT8+, supporting type I TRC identity (**Fig 3F**). KRT8+/NTPDase2^neg^, likely type II and III TRCs, were SOX2-GFP^neg^ (**Fig 3F**), consistent with limited SOX2 expression in type II and III TRCs (Suzuki, 2008). *Shh* was also highly enriched in SOX2^High^ cells (**Fig 3E**). HCR in situ hybridization for *Shh* confirmed colocalization of *Shh* and SOX2-IF in one or two cells in the basal compartment of taste buds (**Fig 3G**). Thus, SOX2^High^ cells include both type I TRCs and SHH+ postmitotic taste precursor cells (Miura et al., 2006; Miura *et al*., 2014).

We next interrogated the dataset for GO term enrichment (**File S2**). With significance at p<0.01, SOX2^High^ cells were enriched for 145 GO terms, SOX2^HiMed^ for 24, and SOX2^Low^ for 73, while SOX2^MedLow^ cells lacked significantly enriched terms. For SOX2^High^, 30/145 top terms were relevant to development, differentiation, or morphogenesis of non-neural tissues, e.g. *lung alveolus development* & *tongue development*. Additional enriched terms were relevant to 1) nervous system development and function (27/145) – *sensory perception of sound & chemical synaptic transmission*, and 2) cell migration/locomotion (14/145) – *neural crest migration & regulation of cell motility*. SOX2^HighMed^ cells were also enriched, but less so, for terms we categorized as development, differentiation, or morphogenesis of non-neural tissues (4/24) – *negative regulation of bone development & branching morphogenesis of an epithelial tube*, with no neural associated terms. Finally, SOX2^Low^ GO terms were primarily relevant to development, differentiation, or morphogenesis of non-neural tissues (34/73) – *regulation of keratinocyte differentiation & hair follicle development*, while nervous system terms, including *olfactory bulb development & axon guidance*, were less frequent (6/73).

Because SOX2^High^ and SOX2^Low^ cells had the most significantly enriched GO terms, we analyzed these further (**File S2**). Terms were grouped into three categories relevant to development, differentiation, or function of: 1) nervous system; 2) soft tissue – kidney, muscle, lung, gut; or 3) hard tissue – bone, teeth, hair, skin. SOX2^High^ were enriched for both neural (27/143) and soft tissue (20/143), with only 2 hard tissue terms, while SOX2^Low^ were enriched for hard tissue (13/73, 7 of top 20) and soft tissue (10/73) GO terms, with only 7/73 related to nervous system.

In sum, neural and endodermal terms were enriched in SOX2^High^ cells, terms relevant to “hard” tissue formation were overrepresented in the SOX2^Low^ population, while intermediate populations (SOX2^HiMed^ and SOX2^MedLow^) were less defined. As TRCs are modified epithelial cells with neural characteristics (Roper, 2007) and non-taste epithelium is keratinized (Cane and Spearman, 1969), GO analysis suggested SOX2^High^ cells have greater taste lineage potential, SOX2^Low^ cells are progenitors of non-taste, keratinized epithelium, but left open the potency of intermediate SOX2-expressing cells. Thus, we next tested if differential SOX2 expression correlates with TRC production in organoids.

### Higher SOX2-expressing cells are taste lineage competent *in vitro*

Lingual organoids were generated from SOX2-GFP+ cells from each fluorescence bin as in **Fig 3A**. qPCR for TRC markers revealed all SOX2 organoids expressed *Kcnj1* (type I marker); *Gnat3, Tas1r2*, and *Pou2f3* (type II); and *Pkd2l1, Snap25*, and *Ascl1* (type III) but at significantly lower levels than LGR5 organoids (**Fig 4A, Fig S1A)**. Among SOX2 organoids, those from SOX2^High^ and SOX2^HiMed^ progenitors tended to have higher TRC marker expression than SOX2^MedLow^ and SOX2^Low^ organoids (**Fig 4A, Fig S1A**). However, whole organoid IF revealed significant differences in TRC differentiation across organoid types. Specifically, SOX2^High^ and SOX2^HiMed^ organoids generated all TRC types – albeit limited production of type III TRCs by SOX2^HiMed^ organoids (**Fig 4B, C**), suggesting taste lineage competency of SOX2^High^ and SOX2^HiMed^ progenitors differ. Further, SOX2^MedLow^ and SOX2^Low^ organoids exhibited little TRC differentiation; organoids generally lacked type III TRCs and had only occasional type I and II TRCs. In sum, although our PCR data suggested some TRC marker expression across all organoid types, IF revealed TRC differentiation was limited primarily to higher expressing SOX2 organoids.

**Figure 4.**
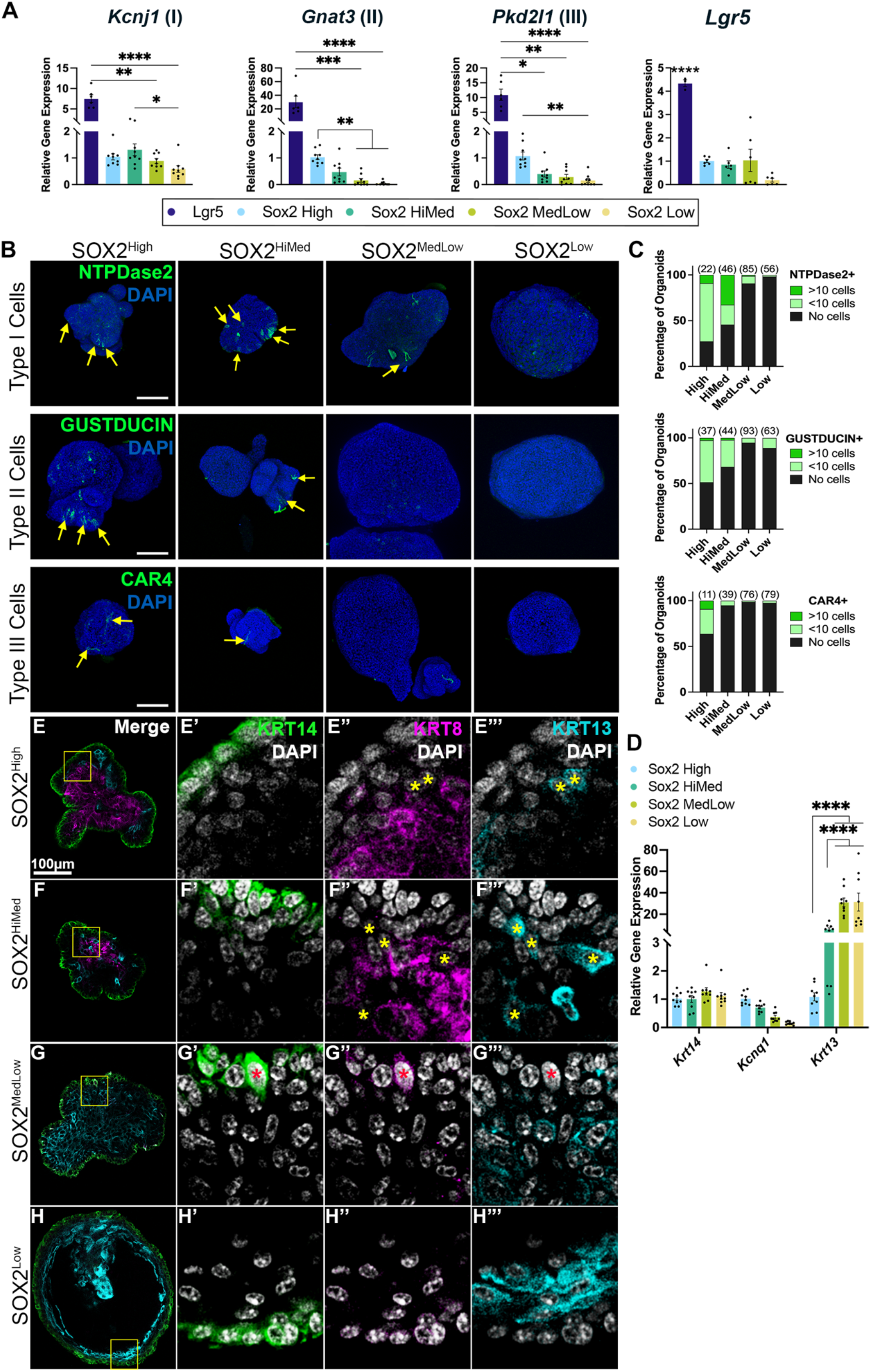
Progenitors with higher SOX2 produce TRC-containing organoids, while organoids from low SOX2 progenitors comprise mostly non-taste epithelium. **A** Expression of type I, II, and III TRC markers (Kcnj1, Gnat3, and Pkd2l1, respectively) trend higher in SOX2^High^ organoids but all SOX2 organoids express these genes at lower levels than LGR5 organoids. **B, C** Most SOX2^High^ and SOX2^HiMed^ organoids contain type I (NTPDase2, green) and type II (GUSTDUCIN, green) TRCs (yellow arrows), and SOX2^High^ organoids contain more type III TRCs than other organoid types (CAR4, green, yellow arrows). Most SOX2^MedLow^ and SOX2^Low^ organoids do not contain TRCs. (n)=total organoids quantified per condition in **C. D** SOX2^High^ and SOX2^HiMed^ organoids express high levels of a general TRC marker, Kcnq1, and moderate Krt13 (non-taste epithelium). SOX2^MedLow^ and SOX2^Low^ organoids express limited Kcnq1 and significantly higher Krt13. Mean ± SEM 1-way ANOVA * p<0.05, ** p<0.01, *** p<0.001 **** p<0.0001. **E-H”** KRT14+ (green) cells make up the external epithelium of all SOX2 organoids (**E’**,**F’**,**G’**,**H’**). SOX2^High^ and SOX2^HiMed^ organoids contain KRT8+ TRCs (magenta **E’’, F’’**) interspersed among KRT13+ non-taste cells (cyan, yellow asterisks, **E’’’**,**F’’’**), while SOX2^MedLow^ and SOX2^Low^ organoids are predominantly KRT13+ cells (cyan **G**,**G’’’**,**H**,**H’’’**). SOX2^MedLow^ organoids have sparse KRT8+ cells that co-express KRT14 (magenta, red asterisk in **G)**. Images in B are maximum projections of confocal z-stack of whole organoids with DAPI nuclear counterstain (blue). Images in E-H”‘ are optical sections of immunostained organoids. Boxed areas in left column are shown in three right columns. DAPI (white) nuclear counterstain.

We also assessed non-taste lineage production in organoids from different SOX2+ progenitors. All SOX2 organoids had comparable *Krt14* expression and KRT14+ progenitors were consistently external (**Fig 4E-H**). As expected, SOX2^High^ and SOX2^HiMed^ organoids expressed *Kcnq1* more highly (**Fig 4D**) and contained more KRT8+ TRCs (**Fig 4E’’, F’’**) than SOX2^MedLow^ and SOX2^Low^ organoids (**Fig 4G’’, H’’**). While SOX2^High^ and SOX2^HiMed^ organoids contained non-taste cells (**Fig 4E’’’, F’’’**), SOX2^MedLow^ and SOX2^Low^ organoids expressed strikingly high levels of *Krt13* (**Fig 4D**) and comprised mostly KRT13+ cells (**Fig 4G’’’, H’’’**). KRT8+ cells were essentially absent in SOX2^Low^ organoids (**Fig 4H’’**); while present sporadically in SOX2^MedLow^ organoids, KRT8+ cells were consistently KRT14+ (**Fig 4G’-G’’’**) reminiscent of immature taste-fated daughters (see above). KRT8+/14+ cells were not observed in SOX2^Low^ organoids suggesting further differences in competency between the two dim SOX2+ populations.

In rodents, non-taste epithelium renews in 5-7 days (Potten et al., 2002), while taste cells renew every 10-30 days (Beidler and Smallman, 1965; Perea-Martinez et al., 2013). SOX2^MedLow^ and SOX2^Low^ organoids, composed of rapidly renewing non-taste epithelium, grew noticeably more than SOX2^High^ and SOX2^HiMed^ organoids, which house taste and non-taste lineages and had growth comparable to LGR5 organoids (**Fig S2A, B**). Compared to other SOX2 populations or LGR5+ cells, SOX2^High^ cells generated ∼50% fewer organoids (**Fig S2C**), consistent with our demonstration that SOX2^High^ cells comprise post-mitotic SHH+ precursors and type I TRCs (see **Fig 3**). Thus, we surmise roughly half of SOX2^High^ cells are taste lineage-competent progenitors.

**Fig S1 for Figure 4.**
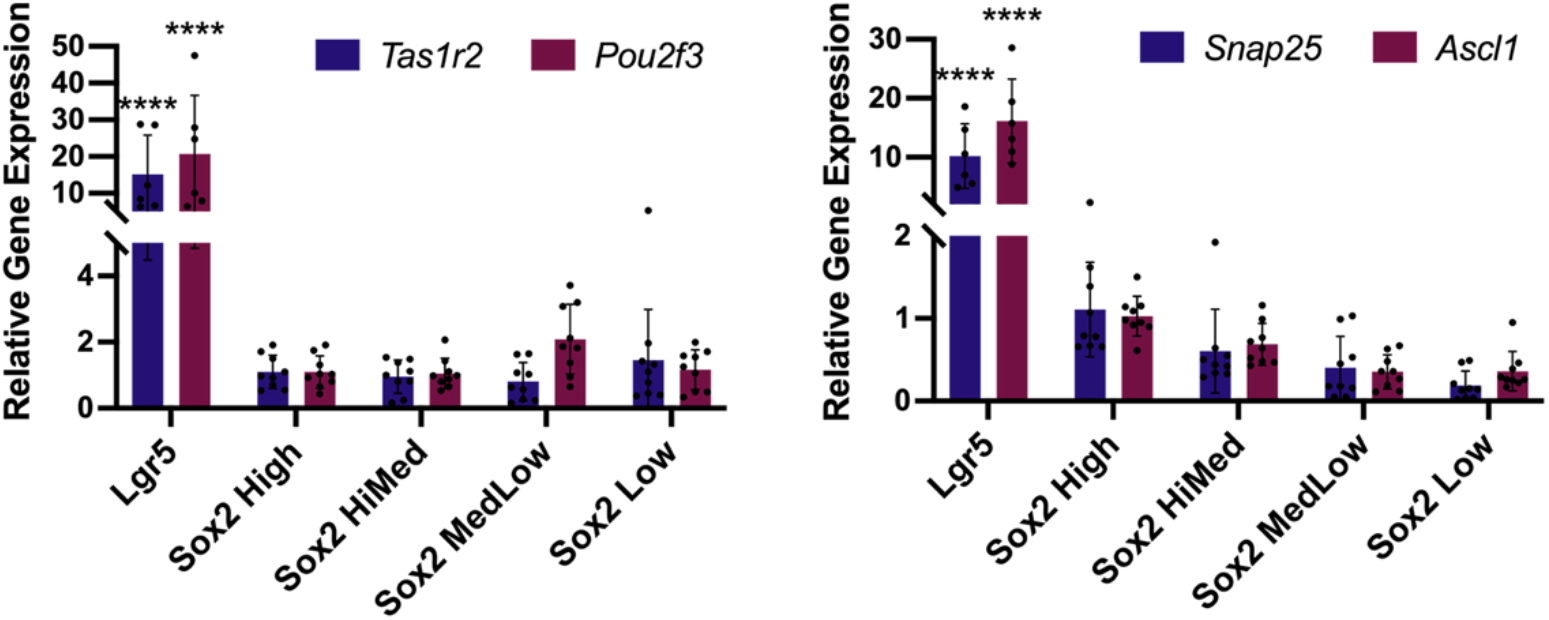
SOX2 progenitor cells generate organoids that express type II and III TRC markers. SOX2 organoids express lower type II (*Tas1r2, Pou2f3*) and type III (*Snap25, Ascl1*) TRC markers than LGR5 organoids. n = 9 biological replicates per organoid population, mean ± S.E.M. 2-way ANOVA **** p<0.0001.

**Fig S2 for Figure 4.**
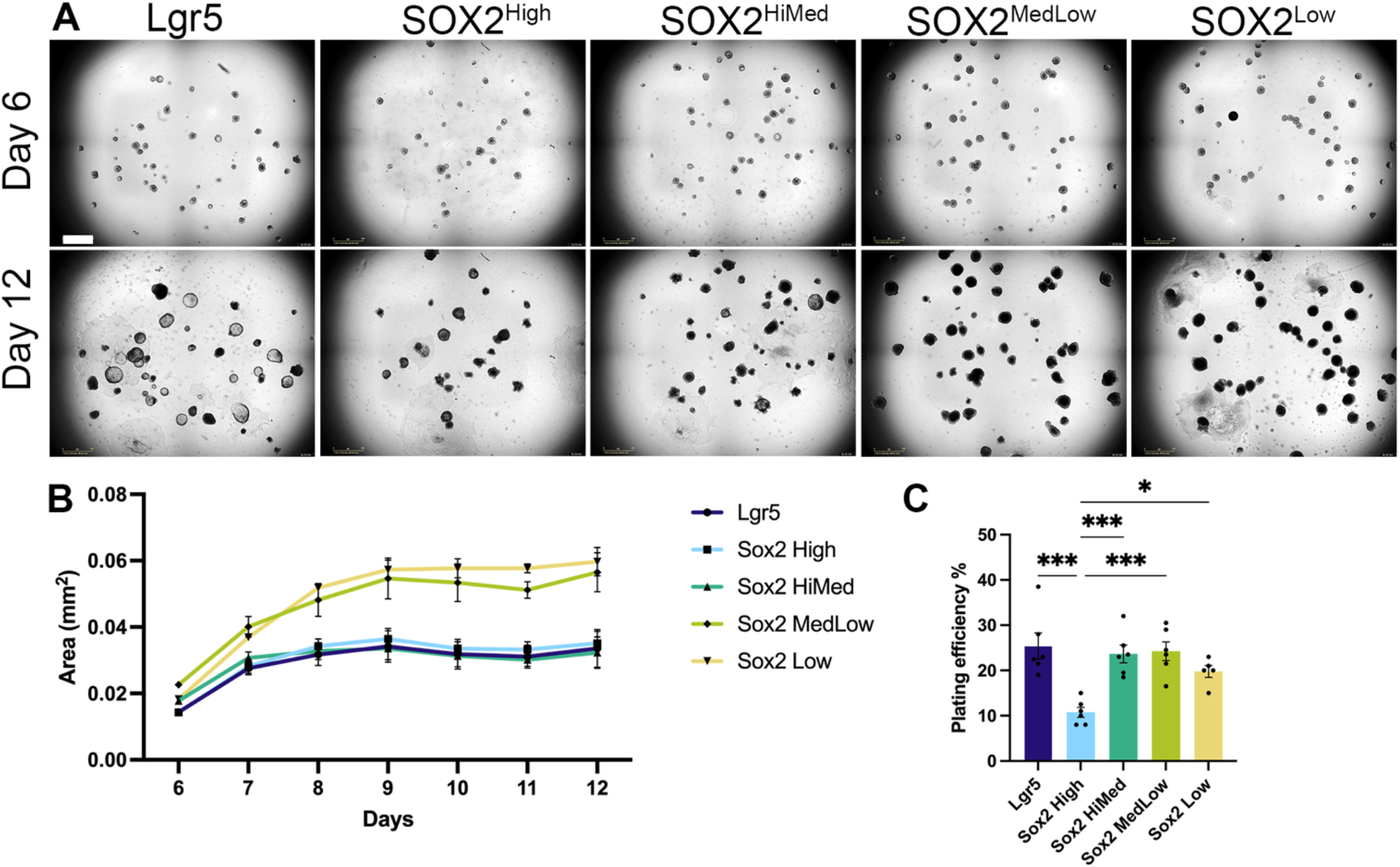
Plating efficiency and growth of organoids from SOX2+ progenitors differ among SOX2 populations. **A** Representative Incucyte images of LGR5 and SOX2 organoids at days 6 and 12. Scale bar, 800µm. **B** SOX2^Medlow^ and SOX2^Low^ organoid growth is more rapid initially and plateaus later than SOX2^High^, SOX2^MedHigh^ and LGR5 organoids. **C** SOX2^High^ cells have a lower plating efficiency than all other progenitors. For **A, B** n = 5-6 experimental replicates per condition. Mean ± SEM 1-way ANOVA * p<0.05, *** p<0.001.

### Hedgehog signaling does not induce TRC differentiation in SOX2+ organoids

*In vivo*, progenitors adjacent to taste buds are sensitive to Hh, i.e., express the Hh target gene *Gli1* (Miura et al., 2001). Transcriptome analysis revealed Hh pathway genes are differentially expressed across SOX2+ populations, including target genes *Gli1* and *Ptch1* (**Fig 5A**). In CVP tissue sections, many SOX2+ basal cells outside of buds are *Gli1*+ (**5B-B”‘**), suggesting some SOX2+ progenitors are Hh-responsive *in vivo*. In fact, Hh is required for TRC differentiation and induces SOX2 expression in adult mouse FFP (Castillo et al., 2014; Castillo-Azofeifa *et al*., 2018). These findings suggested increased Hh signaling could boost taste lineage production in organoids.

**Figure 5.**
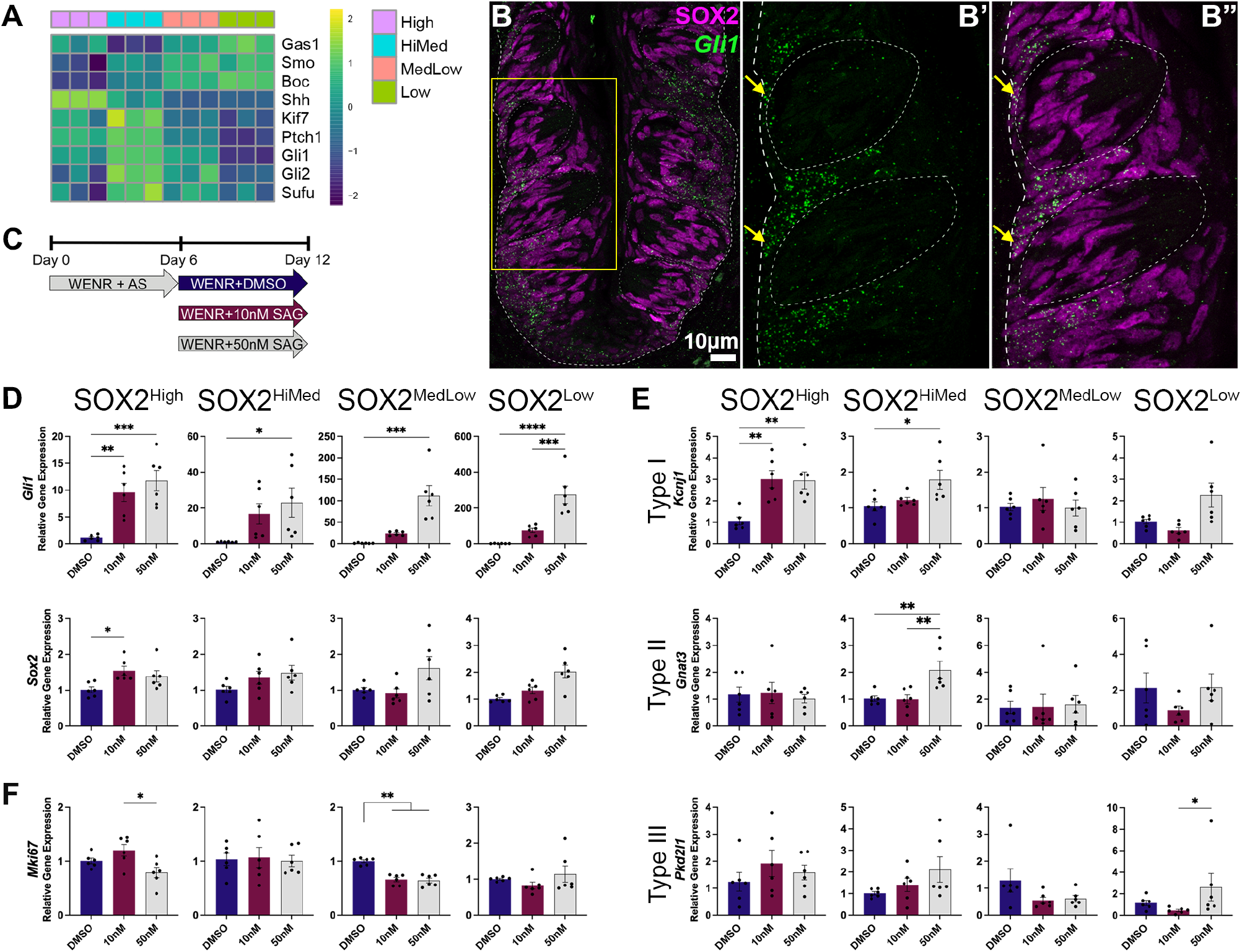
Hedgehog (Hh) pathway activation does not increase TRC differentiation in SOX2 organoids. **A** Hh pathway genes are differentially expressed in SOX2 progenitor populations. **B** *Gli1* (green) is expressed by SOX2-IF cells (magenta) outside of taste buds (dashed circle) (**B’** arrows). Image is a confocal z-stack projection. **C** Organoids were treated with Smoothened agonist (SAG) days 6-12. **D** Hedgehog target gene *Gli1* is significantly upregulated by SAG; *Sox2* is unchanged. **E, F** SAG minimally affects expression of type I (*Kcnj1*), type II (*Gnat3*) and type III (*Pkd2l1*) TRC markers, or proliferation (*Mki67*) in any organoid population. Mean ± SEM 1-way ANOVA, * p<0.05, ** p<0.01.

Organoids from each SOX2+ bin were treated with the Smoothened agonist, SAG, during organoid differentiation (**Fig 5C**). SAG treatment increased *Gli1* expression in all organoid types yet *Sox2* expression was unchanged (**Fig 5D**), suggesting factors required *in vivo* for upregulation of SOX2 by Hh are absent *in vitro*. SAG also had little impact on TRC marker expression; *Kcnj1* (type I) was upregulated in SOX2^High^ and SOX2^HiMed^ organoids, and *Gnat3* (type II) increased in SOX2^HiMed^ organoids (**Fig 5E**). Hh activation did not drive TRC differentiation in SOX2^MedLow^ and SOX2^Low^ organoids, which likely remained mostly non-taste epithelial. Although SAG treatment increased *Pkd2l1* expression (type III) in SOX2^Low^ organoids, no increase was detected in another type III marker, *Ascl1* (**Fig S3**). Our results are consistent with previous reports, where SHH did not impact TRC differentiation in LGR5 organoids (Ren *et al*., 2014). Hh pathway inhibition reduced expression of a proliferation marker, *Mki67*, in LGR5 organoids (Ren et al., 2017), but here SAG did not increase *Mki67* expression in SOX2 organoids (**Fig 5F**). In sum, Hh appears dispensable for progenitor proliferation and TRC differentiation in SOX2-derived organoids.

**Fig S3 for Figure 5.**
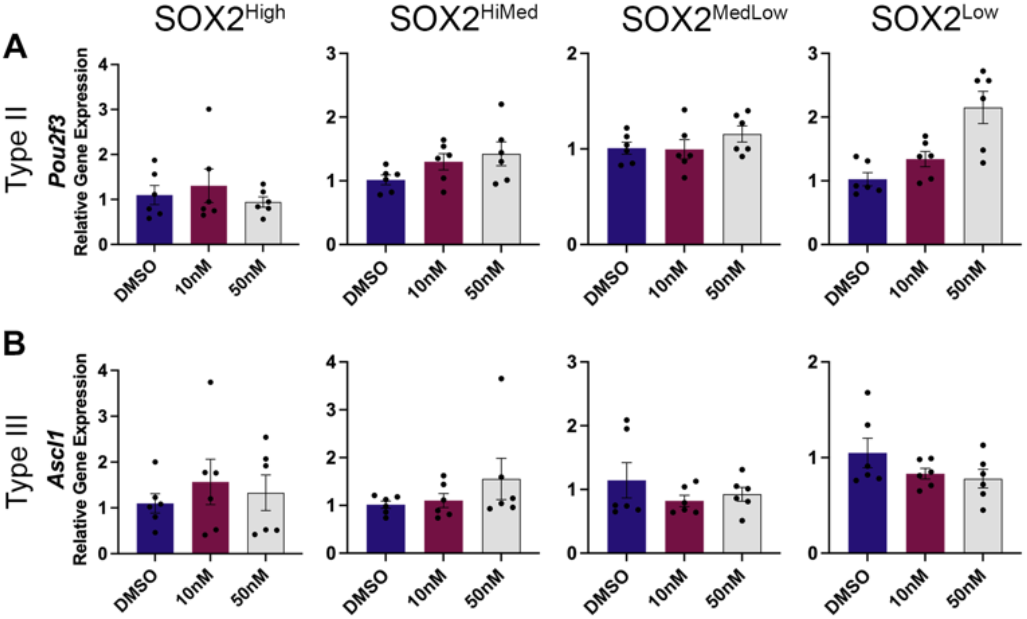
Expression of type II (*Pou2f3*) and III (*Ascl1*) TRC markers is unchanged in SOX2 organoids treated with Smoothened Antagonist (SAG). n = 6 biological replicates/condition. Mean ± SEM.

### Beta-catenin enhances TRC production in organoids of higher expressing SOX2 progenitors

WNT/ß-catenin regulates taste bud homeostasis *in vivo* (Gaillard and Barlow, 2021) and is essential for differentiation of TRCs in LGR5 organoids (Ren *et al*., 2014). WNT pathway transcripts are differentially expressed among SOX2+ cells, with target genes, *Tcf7, Tcf7l1, Lef1, Lgr5* and *Lgr6*, and Frizzled receptors, *Fzd1, Fzd2, Fzd3, Fzd4, Fzd8* and *Fzd10* enriched in SOX2^High^ and SOX2^HiMed^ cells, and ligands, *Wnt3, Wnt3a, Wnt10a*, and *Wnt10b*, enriched in SOX2^MedLow^ and SOX2^Low^ progenitors (**Fig 6A**). Our culture medium includes ample WNT3A, but since Fzds are low in SOX2^MedLow^ and SOX2^Low^ cells, exogenous WNT ligand may be insufficient to drive TRC formation in organoids from these progenitors.

**Figure 6.**
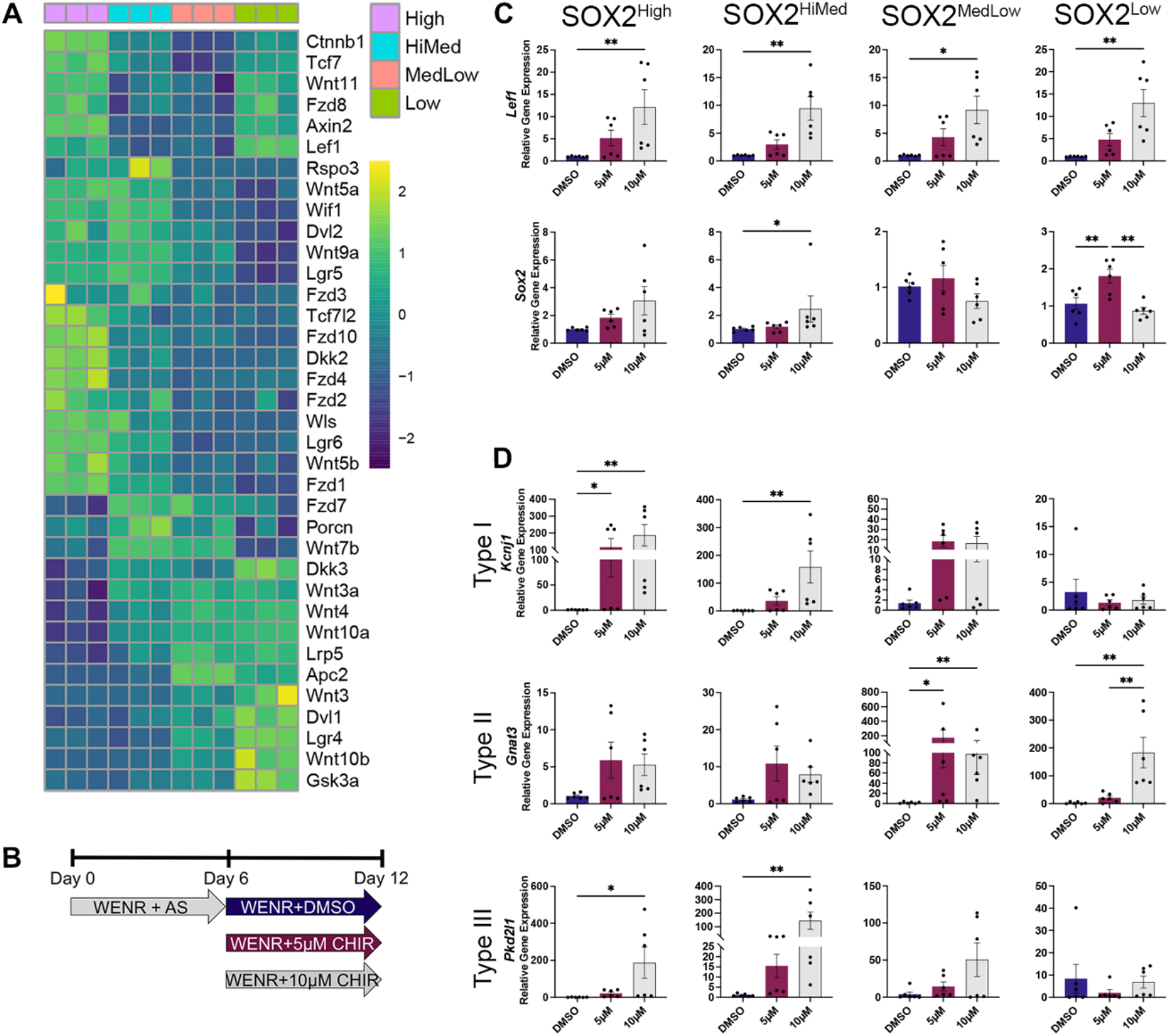
Beta-catenin augmentation enhances TRC production only in organoids from higher expressing SOX2+ progenitors. **A** WNT/ß-catenin signaling pathway genes are differentially expressed across SOX2 populations. **B** Organoids were treated with CHIR99021 (CHIR) from day 6-12. **C** WNT target *Lef1* is significantly upregulated by CHIR in all organoids, but *Sox2* expression is unaltered. **D** In SOX2^High^ and SOX2^HiMed^ organoids, CHIR significantly increases expression of type I (*Kcnj1*) and type III (*Pkd2l1*) TRC markers, while type II TRC marker (*Gnat3*) expression trended higher. In SOX2^MedLow^ organoids, *Kcnj1* and *Pkd2l1* trend upward in response to CHIR, and *Gnat3* was significantly upregulated. In SOX2^Low^ organoids, CHIR also caused elevated *Gnat3*, but had no effect on other TRC markers. Mean ± SEM 1-way ANOVA, * p<0.05, ** p<0.01.

To test if increased ß-catenin downstream of Fzds increased or promoted TRC production in SOX2^High^ and SOX2^HighMed^, or SOX2^MedLow^ and SOX2^Low^ organoids, respectively, cultures were treated with CHIR99021 (CHIR), which upregulates ß-catenin signaling downstream of FZDs (An et al., 2010) (**Fig 6B**). *Lef1*, a WNT/ß-catenin target gene, was significantly increased in all CHIR-treated SOX2 organoids (**Fig 6C**). Although WNT/ß-catenin upregulates SOX2 in embryonic tongues (Okubo *et al*., 2006), CHIR’s effect on *Sox2* expression was inconsistent in organoids (**Fig 6C**). However, TRC marker expression was increased in CHIR-treated SOX2^High^ and SOX2^HiMed^ organoids, consistent with their taste competency (above). Finally, induction of TRC marker expression in SOX2^MedLow^ organoids was highly variable but trended higher, while SOX2^Low^ organoids were largely unaffected by CHIR (**Fig 6D**).

## DISCUSSION

In the CVP of adult mice, TRC renewal is supported by LGR5+ and SOX2+ progenitors (Finger and Barlow, 2021) for review), however, questions remain as to whether LGR5+ and SOX2+ progenitors have comparable taste lineage potential. Further, SOX2 is variably expressed by basal keratinocytes and by a subset of postmitotic cells within taste buds, suggesting SOX2 cells likely represent a mixed cell population, not all of which are taste-competent progenitors.

Individual LGR5+ progenitors generate CVP taste and non-taste lineages in organoids. To test the taste potency of SOX2+ CVP cells we isolated differentially expressing SOX2-GFP+ cells from adult mouse CVP and compared their ability to generate lingual organoids containing both taste and non-taste lineages to that of LGR5+ progenitors. We find higher expressing SOX2+ progenitors are taste competent, and produce organoids composed of all TRC types and non-taste cells. Additionally, our analysis reveals roughly half of SOX2 high cells are type I glial-like TRCs confirming (Suzuki 2008), or SHH+ taste precursor cells (Miura et al., 2014), which are both postmitotic and therefore unlikely to generate organoids. Low expressing SOX2+ progenitors by contrast, are minimally taste-competent and generate organoids composed primarily of non-taste epithelium. Our results are consistent with previous suggestions that high expressing SOX2+ basal keratinocytes are taste progenitors (Castillo-Azofeifa *et al*., 2018; Ohmoto *et al*., 2017; Okubo *et al*., 2006). Importantly, organoid technology allows dissection of the potency of discrete progenitor populations – at least *in vitro*, which is difficult to accomplish with genetic mouse models *in vivo*.

### SOX2 levels allow fine tuning of development and homeostasis

SOX2 dosage affects the behavior of embryonic stem cells (ESCs), as small experimentally induced changes in expression can drive ESC differentiation, e.g. (Kopp et al., 2008). In embryonic neural ectoderm, the levels of SOX2 and other SOXB family members tightly regulate the balance between progenitor proliferation and differentiation, with overexpression driving proliferation and depletion leading to cell cycle exit and differentiation (see (Sarkar and Hochedlinger, 2013). In murine hippocampus, SOX2 is required for stem cell maintenance (Ellis et al., 2004) and in trachea it is required for progenitor proliferation and genesis of the proper proportions of functional cell types (Que *et al*., 2009). The dosage-dependent effects of SOX2 are mediated by interactions with transcriptional cofactors, which themselves are dependent on SOX2 level (Sarkar and Hochedlinger, 2013). In taste epithelium, SOX2 is required for continual renewal of TRCs in FFP and CVP (Castillo-Azofeifa *et al*., 2018; Ohmoto et al., 2020; Okubo *et al*., 2006), and higher SOX2 expression is associated with progenitors adjacent to taste buds in each taste field (Castillo-Azofeifa *et al*., 2018; Okubo *et al*., 2006). Here our organoid data suggest differential SOX2 expression predicts the taste lineage potential of CVP progenitors, however, the transcriptional cofactors that affect SOX2 function remain to be explored.

### Molecular regulation of taste bud renewal in organoids

In adult mice, Hedgehog (Hh) and WNT/ß-catenin regulate taste epithelial homeostasis. Hh is required for taste bud maintenance as inhibition abolishes pro-TRC differentiation signals (Castillo-Azofeifa et al., 2017; Castillo-Azofeifa *et al*., 2018; Lu et al., 2018). Hh functions upstream of SOX2, as overexpression of SHH in lingual progenitors induces excess and ectopic taste buds that express high SOX2 (Castillo *et al*., 2014; Golden et al., 2021); SOX2 is also required for the formation of endogenous and ectopic taste buds downstream of Hh (Castillo-Azofeifa *et al*., 2018; Okubo *et al*., 2006). In lingual organoids derived from differentially expressing SOX2+ progenitors, SAG treatment increased Hh target gene expression (*Gli1*), but *Sox2* expression was not induced nor was TRC differentiation augmented. We have argued that Hh may function *in vivo* to promote changes in cell adhesion and locomotion permissive to TRC differentiation from progenitors located outside of taste buds, processes that are in part regulated by SOX2 (Golden *et al*., 2021). However, since taste bud structures do not form in organoids, we hypothesize Hh and its regulation of SOX2 are dispensable *in vitro*.

In mice, WNT/ß-catenin is required for progenitor proliferation and survival, and at higher levels, impacts TRC fate (Gaillard and Barlow, 2021). WNT is also required for many epithelial organoid systems including lingual organoids to support growth (Ren *et al*., 2014; Shechtman *et al*., 2021). Transcriptome analysis revealed variable WNT pathway gene expression in different SOX2+ populations, suggesting TRC production from lower expressing SOX2+ cells might require pathway augmentation. However, pharmacologically increased WNT/ß-catenin enhanced TRC production only in taste competent SOX2-bright but not SOX2-dim organoids, suggesting not all progenitors are competent to respond to increased WNT signaling with TRC production. These observations are consistent with findings *in vivo* in anterior tongue, where genetic induction of stabilized ß-catenin in lingual progenitors throughout the tongue increases differentiation of TRCs but only in taste bud-bearing FFP and not elsewhere in tongue epithelium (Gaillard *et al*., 2015). Notably, SOX2 expression is robust in FFP epithelium (Castillo-Azofeifa *et al*., 2018) where additional TRCs are induced by increased WNT/ß-catenin, and very low in non-taste epithelium where elevated WNT/ß-catenin does not drive excess TRC production, suggesting SOX2 level predicts taste competency in vivo as well as in organoids.

In summary, lingual organoid technology provides a rapid means of testing the competency of mouse taste stem cells, including pharmacological manipulation and gene profiling. This in vitro approach can complement and extend findings *in vivo*, and will be an important tool to explore taste epithelial homeostasis.

## METHODS

Mice were obtained from Jackson Laboratory (*Lgr5*^*EGFP-IRES-CreERT2*^ #008875; *Sox2*^*GFP*^ #017592) and maintained in an AAALAC-accredited facility in compliance with the Guide for Care and Use of Laboratory Animals, Animal Welfare Act and Public Health Service Policy. Procedures were approved by the Institutional Animal Care and Use Committee at the University of Colorado Anschutz Medical Campus.

### Organoid production

was as described (Shechtman *et al*., 2021). Tongue epithelium from 4-8 *Lgr5*^*EGFP*^ or *Sox2*^*EGFP*^ mice aged 8-20 weeks were used per experiment. Briefly, Collagenase (2mg/mL) and Dispase (5mg/mL) in PBS was injected beneath and beside the CVP, the epithelium peeled and then dissociated for 45 min in Collagenase (2mg/mL), Dispase (5mg/mL), and Elastase (2mg/mL) at 37°C. Cells were centrifuged (2000 RPM 4°C), the pellet resuspended in FACS buffer (1mM EDTA, 25mM HEPES pH 7.0, 1% FBS, 1x Ca^2+^/Mg^2+^-free dPBS), passed through a 30µm cell strainer and subjected to FACS for GFP signal. GFP^neg^ cells were discarded. Sorted GFP+ cells were plated in 48-well plates at 200 cells/well in 15µL Matrigel in WENR: 50% WRN conditioned media (Miyoshi and Stappenbeck, 2013); 1X Glutamax, 1X HEPES, 1X penicillin-streptomycin, 1X B27 Supplement, 1X Gentamicin (<u>Gibco</u>); 1X Primocin (<u>InvivoGen)</u>; 25ng/mL Murine Noggin, 50ng/mL Murine EGF (<u>Peprotech)</u>; 1mM Nicotinamide, 1mM N-acetyl-L-cysteine (<u>Sigma</u>); 10µM Y27632 (<u>Stemgent</u>) was used days 0-2 to promote survival, and 500nM A8301 (<u>Sigma</u>) and 0.4µM SB202190 (<u>R&D Systems</u>) for growth (days 0-6). Medium was changed every 2 days.

### Tissue collection for IF and HCR

*Lgr5*^*EGFP*^ and *Sox2*^*GFP*^ mice were perfused transcardially with periodate-lysine-paraformaldehyde, tongues dissected from the jaw, incubated for 3h at 4°C in 4% paraformaldehyde (PFA) in 0.1M phosphate buffer (PB) and placed in sucrose (20% in 0.1M PB) overnight at 4°C. Samples were embedded in OCT Compound (Tissue-Tek), 12µm cryosections collected on SuperFrost Plus slides (Fisher) and stored at -80°C.

### Immunofluorescence

Sections were washed in 0.1M phosphate buffered saline (PBS), incubated in blocking solution (BS: 5% normal goat or donkey serum, 1% bovine serum albumin, 0.3% Triton X100 in 0.1M PBS, pH 7.3) for 1.5h at room temperature (RT) followed by primary antibodies diluted in BS for 2 nights at 4°C. Sections were rinsed in PBS + 0.1% Triton, incubated for 1h, RT in secondary antibodies in BS, followed by DAPI nuclear counterstain. Slides were coverslipped with ProLong Gold (Thermo Fisher). Organoids were harvested as described (Shechtman *et al*., 2021). Briefly, organoids were incubated in Cell Recovery Solution (Corning) at 4°C, washed in 0.1M PBS, fixed in 4% PFA, and stored at 4°C in PBS with 1% BSA. For IF, organoids were incubated in BS (2h), then with primary antibodies in BS for 3 nights at 4°C, washed with PBS + 0.2% Triton, and incubated with secondary antibodies in BS overnight at 4°C. Organoid nuclei were counterstained with DAPI, washed with 0.1M PB and mounted on SuperFrost Plus slides in Fluoromount (Southern Biotech). For dispersed cell immunostaining, CVP epithelia from 2 *Sox2*^*GFP*^ mice were dissociated, and cells mounted on poly-D-lysine (1:10 in H2O) and Fibronectin (1:100 in 1x PBS) coated coverslips, fixed in 4% PFA for 2 min RT, washed with 0.1M PBS, and stored at -80°C until processed for IF as for tissue sections. Antibodies are listed in **Table 1**.

**Table 1.**
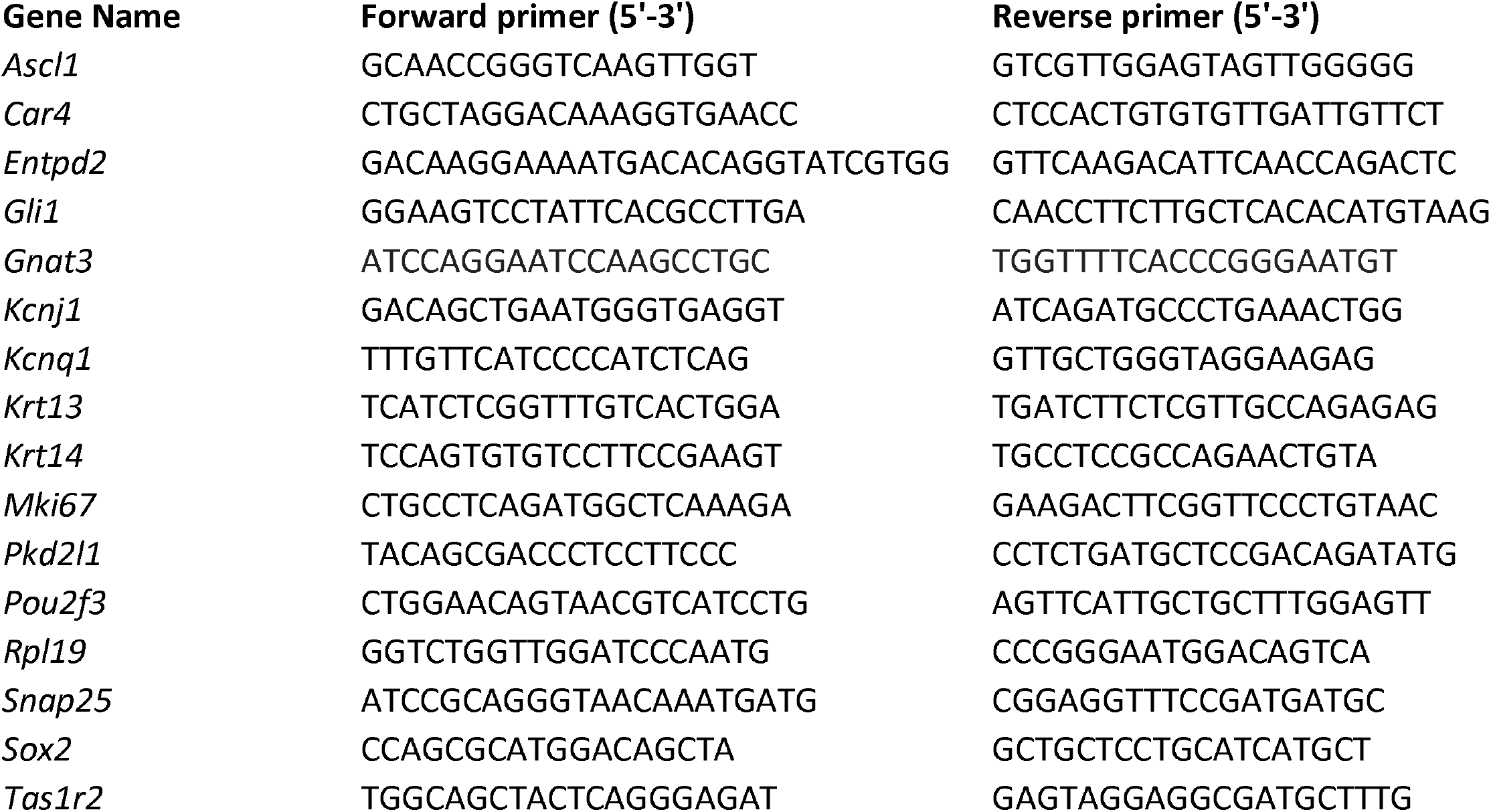
List of qPCR primer sequences and antibodies.

**Table 2.** Summary of GO analysis.

| **Antibodies** | **Catalog Number** | **Dilution** |
| --- | --- | --- |
| DAPI (for FACS) | Thermo Fischer 62247 |  |
| DAPI (for IHC) | Invitrogen D3571 | 1:10,000 |
| Chicken anti-GFP | Aves Labs; GFP-1020 | 1:250 |
| Goat anti-CAR4 | R&D Systems; AF2414 | 1:50 |
| Guinea pig anti-KRT13 | Acris Antibodies; BP5076 | 1:250 |
| Rabbit anti-GUSTDUCIN | Santa Cruz Biotechnology; sc-395 | 1:250 |
| Rabbit anti-KRT14 | BioLegend; 905301 | 1:1000 |
| Rabbit anti-NTPDase2 | CHUQ; mN2-36LI6 | 1:300 |
| Rabbit anti-SOX2 | Cell Signaling; 23064S | 1:100 |
| Rat anti-KRT8 | DSHB;TROMA-IS | 1:100 |

### HCR

Molecular Instruments designed and produced probes against *Sox2* (NM_011443.4), *Lgr5* (NM_010195.2), and *Shh* (NM_009170.3). Methods were adapted from the manufacturer’s protocol. Frozen sections were incubated in 4% PFA for 10 min, RT, washed with 0.1M PBS, and incubated in 2 µg/mL Proteinase K for 2 min, RT. Sections were incubated in Triethanolamine Solution, 12M HCl, and Acetic anhydride in DEPC water for 10 min RT, washed with PBS, incubated in hybridization buffer for 55 min at 37°C, and then in 1.2 pmol probe in hybridization buffer overnight at 37°C. Sections were washed with: 75% wash buffer/25% 5x SSCT (20x SSC, 10% Tween20, ultrapure water), 50% wash buffer/50% 5x SSCT, 25% wash buffer/75% 5x SSCT, and 100% 5x SSCT, each for 20 min at 37°C, then 100% 5x SSCT for 20 min, RT. Sections were incubated in amplification buffer for 1h, then in denatured hairpin solution (6 pmol hairpin 1 and 2 in amplification buffer) overnight. Slides were washed with 5x SSCT and mounted in DAPI-containing ProLong Gold.

### Image acquisition and analysis

CVP sections and organoids were imaged with a Leica TCS SP8 laser-scanning confocal microscope with LAS X software. Sequential z-stacks of CVP sections were acquired as 0.75 µm optical sections, and organoids via 2 µm optical sections. For analysis, investigators were blind to condition. Immunolabeled cells per organoid were tallied when: 1) a cell is immunomarker-positive; and 2) an immunostained cell has a DAPI+ nucleus.

### Quantitative RT-PCR

Organoids were harvested as described (Shechtman *et al*., 2021). Briefly, plates were placed on ice for 30 min and organoids freed from matrix by scratching with a pipet tip. Three wells were pooled per biological replicate, organoid samples were centrifuged and resuspended in RLT buffer (Qiagen) with ß-mercaptoethanol. RNA was extracted via RNeasy Micro Kit (Qiagen), quantified via Nanodrop (ThermoFisher Scientific), and reverse transcribed with an iScript cDNA synthesis kit (Bio-Rad). Power SYBR Green PCR Master Mix (Applied Biosystems) was used for qPCR reactions on a StepOne Plus Real-Time PCR System (Applied Biosystems, Life Technologies). Relative gene expression was assessed via the ΔΔCT method, with *Rpl19* as the housekeeping gene. Primers are listed in **Table 1**.

### Growth curve and plating efficiency analysis

Growth curves were generated from Incucyte® Live Cell Analysis System (Sartorius) 2-D images of organoids taken daily from days 6-12. ImageJ “analyze particles tool” was used to 1) set the range for expected organoid size (0.003-0.4) mm^2^ and 2) obtain areas of organoids within this range. Organoids were then manually reviewed and non-organoids outside the range criteria excluded from analysis. To address the problem >1 spatially overlapping organoids, these were outlined and individual area measurements taken manually. If manual delineation was not possible these organoids were excluded from area measurements but included in calculating plating efficiency. After manual review, total organoid number identified in each well at day 12 was divided by the starting number of cells (200/well) to obtain plating efficiency.

### RNA Sequencing

CVP epithelia from 11-12 week old *Sox2*^*GFP*^ mice (7-13 mice/replicate) were dissociated and GFP+ cells isolated via FACS as above. RNA was extracted with an Arcturus PicoPure RNA Isolation Kit (ThermoFisher) and stored at -80°C. Poly-A selected sequencing libraries were prepared using Nugen Universal Plus mRNA kit and sequenced (2×150bp) using an Illumina NOVASeq6000 by the Genomics and Microarray Core Facility at the University of Colorado AMC. Reads were trimmed and filtered using BBDuk (v38.50) https://sourceforge.net/projects/bbmap/, resulting in an average of 49.6M (+/-stdev 8.8M) read pairs per sample. Transcript abundance was quantified using Salmon (v1.4.0) (Patro et al., 2017) and a decoy-aware transcriptome index (Gencode M26) (Frankish et al., 2021) at a mapping rate of 85 % (+/-stdev 2.5). Transcript abundance was summarized at the gene-level using ‘tximport’ (Soneson et al., 2015) and differential expression was tested with DESeq2 (Love et al., 2014). A likelihood ratio test was performed to identify differentially expressed genes (DEGs) across all 4 samples. To identify DEGs specific to each SOX2GFP bin, differential expression was tested on each group vs the remaining three. DEG lists from these analyses were used as input to TopGO for gene ontology analysis, with inclusion criteria of average normalized expression across all bins >100, log2FoldChange >1, padj >0.05. A final inclusion criterion included padj >0.05 by the above-mentioned likelihood ratio test analysis. Heatmaps were generated using ‘pheatmap’ https://CRAN.R-project.org/package=pheatmap in R using default column and row clustering methods. Display values are relative expression levels.

### Statistical analyses

Normally distributed data were analyzed by ordinary one-or two-way ANOVA with Tukey’s multiple comparisons post hoc test using GraphPad Prism software. Dunn’s multiple comparison test was used when data were not normally distributed. Data are represented as mean +/-SEM, and significance was taken as p<0.05 with a confidence interval of 95%.

---

**Supplementary File 1. Differential expression analysis across SOX2-GFP progenitors using a likelihood ratio test**. Tab **A** lists all genes. Tab **B** lists genes with ‘padj’ < 0.05. For each tab, columns C-D are average normalized counts per bin. Columns G and H are average normalized counts for bins producing taste-replete organoids (SOX2^High^, SOX2^HiMed^), and those producing non-taste organoids (SOX2^MedLow^, SOX2^Low^). Column I is the False Discovery Rate adjusted p value.

**Supplementary File 2. RNA sequencing analysis of SOX2+ progenitors from adult mouse CVP. A-D** List of differentially expressed genes in SOX2^High^, SOX2^HiMed^, SOX2^MedLow^, and SOX2^Low^ cells. Analysis was performed by comparing each individual bin versus the other three. Only genes with log2FoldChange >0, and FDR-adjusted p < 0.05 are reported. Columns C-D show the average normalized counts per bin. Columns G and H show average normalized counts for bins producing taste-replete organoids (SOX2^High^, SOX2^HiMed^), and those producing primarily non-taste organoids (SOX2^MedLow^, SOX2^Low^). **E-H** List and analysis of Gene Ontology (GO) terms associated with genes enriched in SOX2^High^, SOX2^HiMed^, SOX2^MedLow^, and SOX2^Low^ cells. **I, J** Highlighted list of GO terms associated with development and differentiation of the nervous system, hard tissues, or soft tissues

