## Supplementary figures and images for "High Sox2 expression predicts taste lineage competency of lingual progenitors in vitro"

### Supplemental fig 1

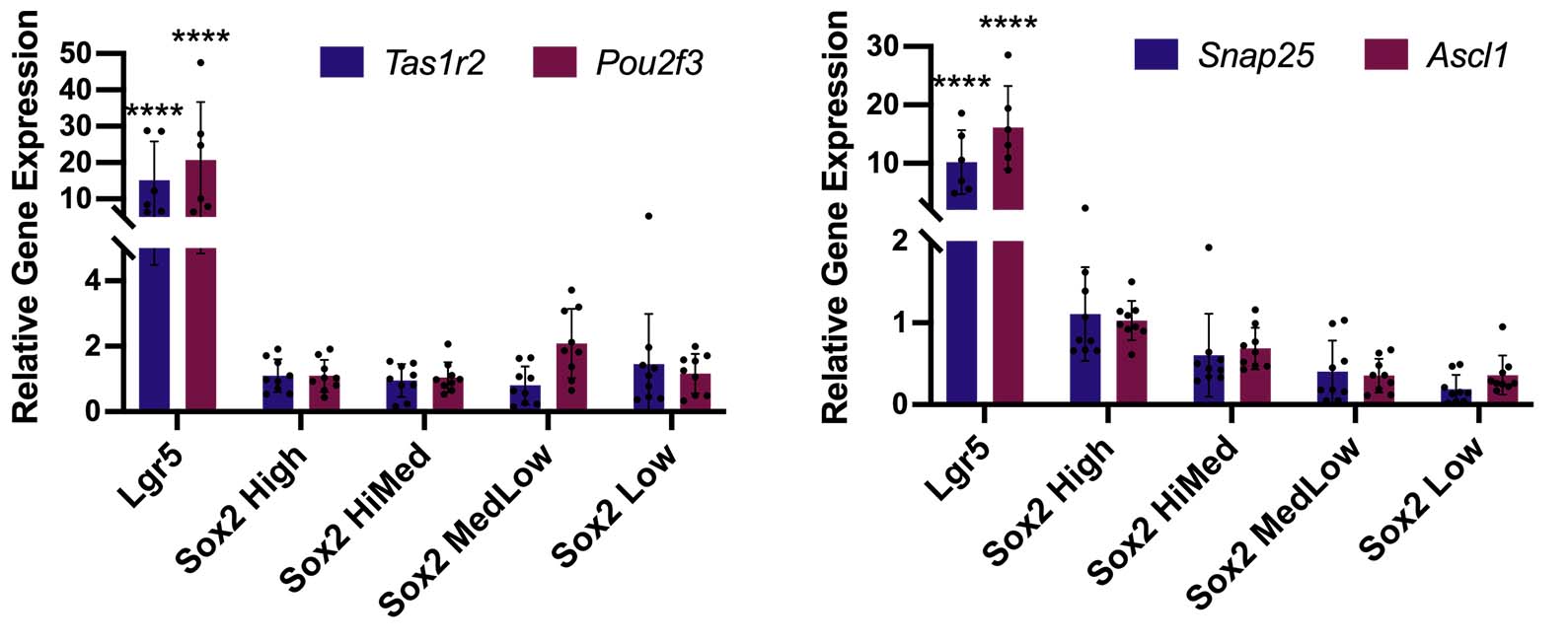

### Supplemental fig 2

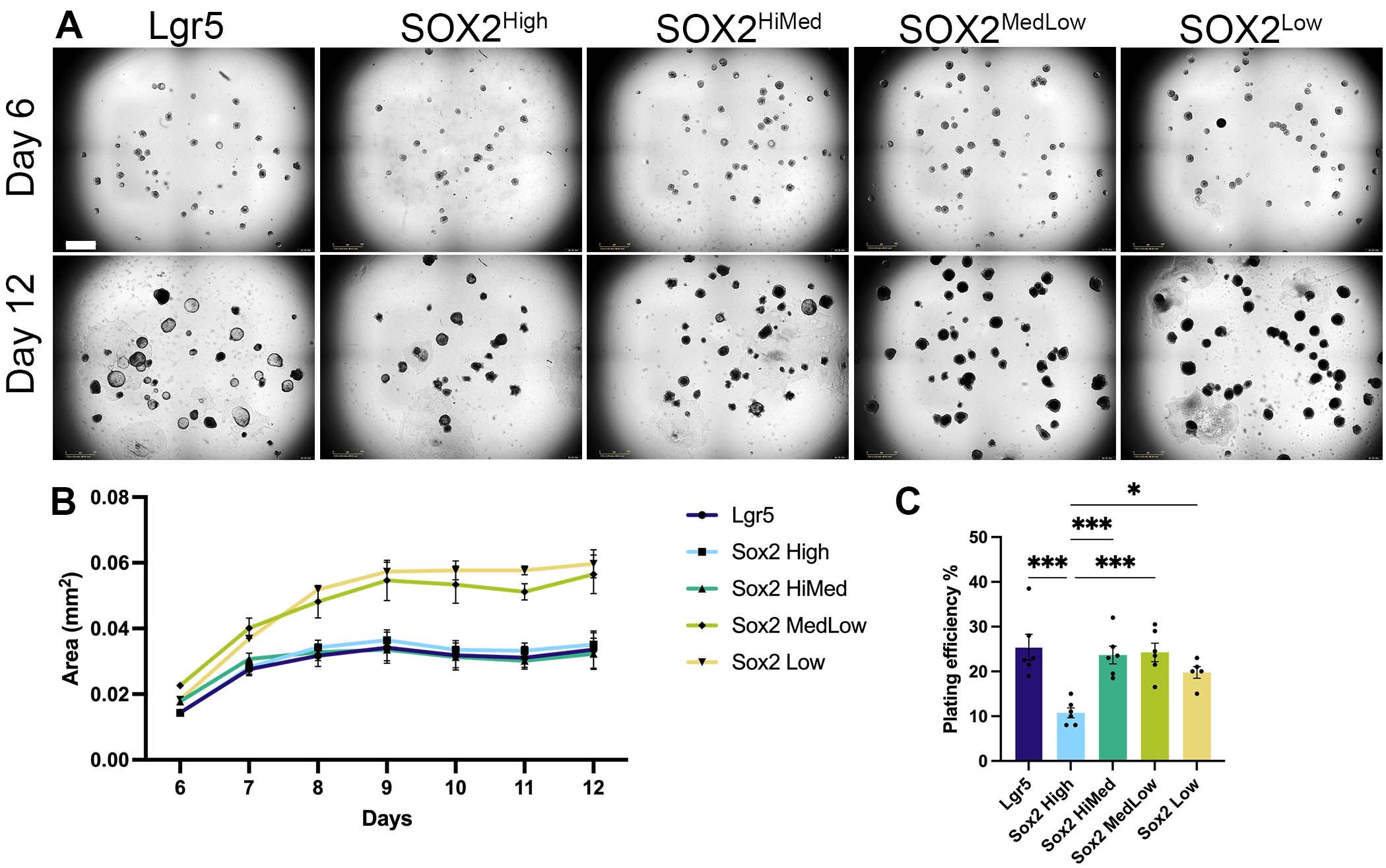

### Supplemental fig 3

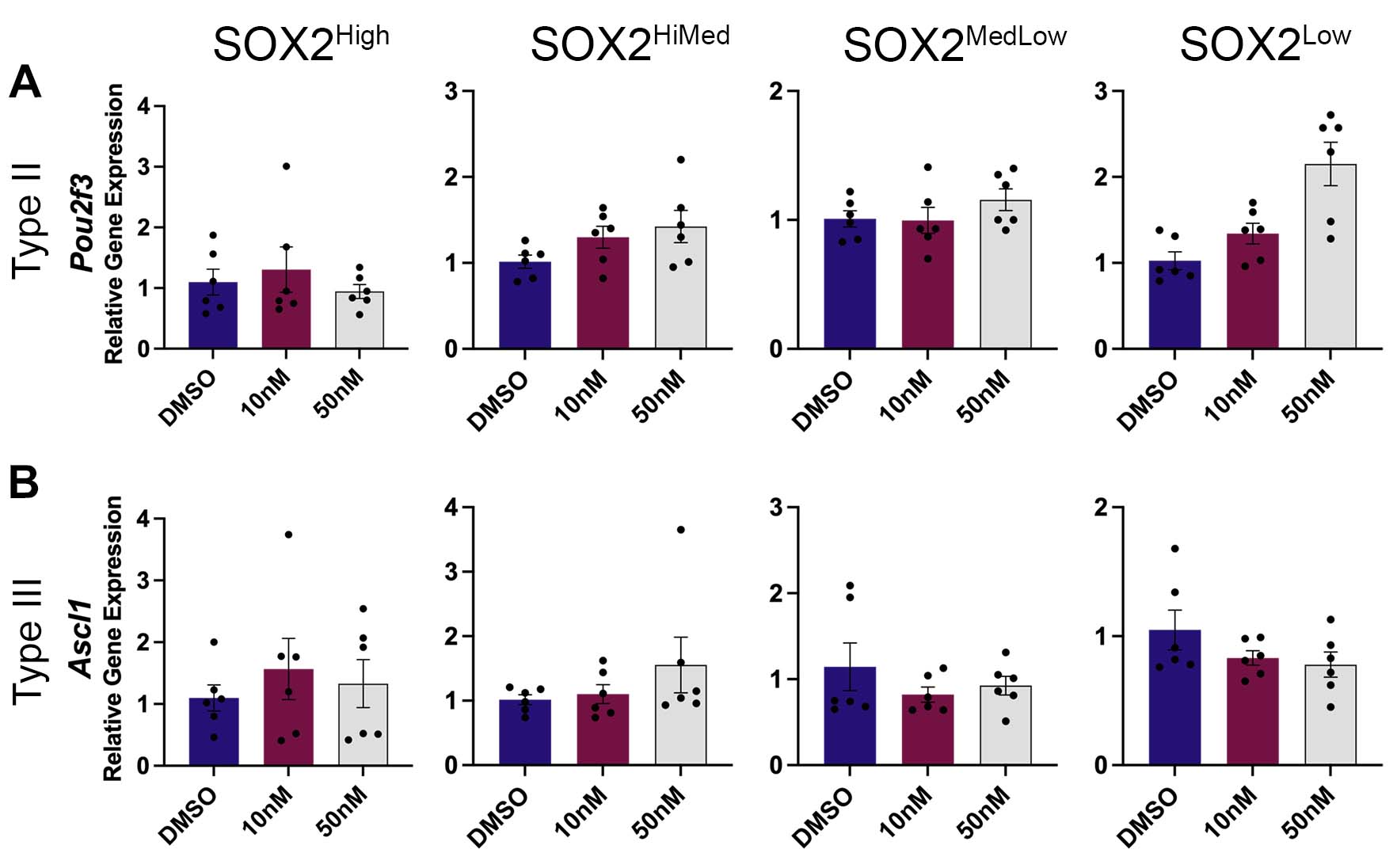
